# Evaluation of microbial diversification mechanisms in legume-based mixed cropping systems with different legume species and types of fertilizer management

**DOI:** 10.1101/2020.12.09.416966

**Authors:** Akari Kimura, Yoshitaka Uchida

## Abstract

Biodiversity loss is becoming a global concern due to its negative impact on services associated with the ecosystem. For agricultural soil to maintain these multi-services, the conservation of soil microbial diversity is of utmost importance. Mixed cropping systems involve the utilisation of multiple crop species on the field as well as the diversification of aboveground plants, although several contradicting results have been reported regarding their impacts on soil microbial diversity. Therefore, for the evaluation of the impact of different leguminous species used in mixed cropping systems as well as types of fertilizer on the diversity of soil microbes, a pot study was performed under maize/legume mixed cropping systems with one of three legumes, including cowpea (*Vigna unguiculate* (L.) Walp.), velvet bean (*Mucuna pruriens* (L.) DC.), and common bean (*Phaseolus vulgaris* L.) as well as one of three types of fertilizer treatments, namely chemical fertilizer (CF), carbonised chicken manure (CM), or the lack of fertilizer (Ctr). 16S rRNA analyses were conducted using the soils sampled from each pot for soil bacterial diversity assessment. Concerning the results, a decrease in the microbial diversity after CM application was shown by the soil with velvet bean + maize (MM) when compared to the Ctr treatment, while an increase in the microbial diversity was shown by the soil with common bean + maize (PM) under the same condition. In case of the CM application, the abundance of treatment-unique bacteria increased in the PM treatment, although their decrease was observed for the MM treatment. In contrast, the abundance of dominant microbes, including Thaumarchaeota was significantly lower in PM but higher in MM when the CM was applied. Legume species-dependent factors, including nutrient absorption and root exudate composition might be important concerning soil bacterial diversities. For the conservation of soil microbial diversity with mixed cropping, the interaction effect of legume species and fertilizer type should be considered in future studies.

## 1 Introduction

Agricultural production needs to be increased with the growth of the world population and more than 50%-80% increase in food production will be needed by 2050 (Dawson et al., 2016; Keating et al., 2014; Tilman et al., 2011). However, the expansion of agricultural lands has been the primary cause of biodiversity loss in terrestrial ecosystems (Kehoe et al., 2017; Zabel et al., 2019). Particularly, modern agricultural practices, including monoculture cropping systems and the intensive use of inorganic fertilizers and pesticides lead to the degradation of soil as well as the threatening of the land with the loss of genetic diversity (Díaz et al., 2006; Rohr et al., 2019). Therefore, the establishment of efficient agricultural crop production systems compatible with biodiversity conservation is a global challenge for future food security.

Among different cases of biodiversity, the diversity of soil microorganisms is especially important for the stability of agricultural ecosystems as they can be considered the main drivers of biogeochemical reactions that are beneficial for soil health and crop productivity (Chaparro et al., 2012). For example, their involvement is notable in biogeochemical processes that are essential for plant health and growth, including nutrient absorption, immune function, pathogen prevention and stress tolerance (Loreau et al., 2001; Nannipieri et al., 2003). Thus, agricultural management systems to maintain or increase soil microbial diversity must be established.

Among the various agricultural practices that can potentially diversify soil microbes, the use of mixed cropping systems and organic resources have been receiving heightened attention. Legume-based intercropping systems were stated by previous reports to enhance soil microbial diversity and bacterial functions, including carbon (C) fixation and citrate cycle (Gao et al., 2010; Lian et al., 2019; Yu et al., 2020). Moreover, mixed cropping systems with field pea varieties were demonstrated by Horner et al. (2019) to be able to build stronger and larger co-occurrence networks in rhizosphere bacterial communities. These studies suggested that the diversification of crops, including mixed cropping systems, is important to sustain soil microbial diversity and the multifunctionality of the agricultural ecosystem.

In addition to crop diversification, the use of organic amendments, including the application of charcoal produced from organic waste into soil can enhance microbial biomass and microbial diversity (Chen et al., 2016; Francioli et al., 2016; Gong et al., 2009; Gul et al., 2015; He et al., 2008). For example, the combined use of charcoal composed of plant residue and legume-based intercropping systems could effectively increase crop production and enhance the abundance of N-fixing and Phosphorus (P)-solubilizing bacteria in the rhizosphere (Duchene et al., 2017; Liao et al., 2019; Liu et al., 2017). Therefore, the combined use of mixed cropping systems and organic fertilizer might be important for the maintenance of soil microbial diversity and efficient nutrient cycles.

However, complex interactions were shown by previous studies between plant species and different types of fertilization in intercropping systems in association with soil microbial diversity. The response of different intercropping systems to fertilizers was reported to be variable as the nutrient uptake ability is controlled by intercropped plant species (Ghosh et al., 2009; Mahmoud, 2007). This suggests that availability of various nutrients in the soil, one of the factors controlling soil microbial diversity, is influenced by the combination of different fertilizers and plant species used in mixed cropping systems. Thus, the impact of mixed cropping on soil microbial diversity to be possibly influenced by the interaction effects between crop species and fertilizer treatments was assumed.

As a conclusion, studies are needed to examine the effect of interaction between various types of fertilizers and legume species on the microbial community and diversity. To evaluate the mechanisms of soil microbial diversification, studies focusing on ‘rare’ microbial taxa are notably important. It was suggested by recent studies that rare microbial taxa (∼1%–3% within the relative abundance), which appear only under certain fertilization management conditions, take critical role in regard of soil multifunctionality in case of agricultural soils (Chen et al., 2020; Hol et al., 2010; Kurm et al., 2017). This can be due to over-proportional roles played by these rare taxa in soil nutrient cycles, including N and C cycles as well as the degradation of complex chemicals (Jousset et al., 2017). Therefore, for the maintenance of soil multifunctionality, the ecology of rare microbes should be considered together with the microbial diversification process.

In this study, the importance of the effect of interaction between legume species and fertilization for microbial diversification mechanisms in association with a mixed cropping system of maize and legume was hypothesized. Treatment-unique bacterial communities under specific combinations of legume species and fertilizer that might contribute to the improvement of the soil microbial diversity were aimed to be identified in this research. A greenhouse experiment of legume-maize mixed cropping was performed using three legume varieties and chemical or organic fertilizer (carbonized chicken manure) to measure bacterial diversity and community structure based on 16S rRNA analysis.

## 2 Materials and methods

### 2.1 Soil sampling

The soil used in this experiment was sampled from bare land at the university farm located at the Field Science Center for Northern Biosphere, Hokkaido University, Japan (43° 04’ N, 141° 20’ E). The properties of the soil were given in Table 1. The soil type was clay loam with 44.6% sand, 21.5% silt, and 33.9% clay.

**Table 1.** The chemical properties of soil and carbonized chicken manure (CM). The number of replications was three for each property measurement.

| Chemical properties | Soil | CM |
| --- | --- | --- |
| Water content (%) | 19.5 ± 0.1 | 19.1 ± 1.2 |
| C/N | 14.7 ± 0.2 | 10.0 ± 0.4 |
| pH (1:10) | 6.3 ± 0.0 | 10.4 ± 0.3 |
| TN (%) | 0.25 ± 0.01 | 4.06 ± 0.45 |
| TC (%) | 3.9 ± 0.1 | 40.5 ± 3.4 |
| NO<sub>3</sub><sup>-</sup>-N mg kg<sup>-1</sup> | 37.4 ± 0.8 | 6.4 ± 0.6 |
| NH<sub>4</sub><sup>+</sup>-N mg kg<sup>-1</sup> | 1.5 ± 0.5 | 159.3 ± 1.1 |
| P<sub>2</sub>O<sub>5</sub> g kg<sup>-1</sup> | 0.48 ± 0.01 | 84.30 ± 2.45 |
| CaO g kg<sup>-1</sup> | 8.29 ± 0.45 | 187.30 ± 6.85 |
| MgO g kg<sup>-1</sup> | 0.89 ± 0.01 | 24.98 ± 0.44 |
| K<sub>2</sub>O g kg<sup>-1</sup> | 0.67 ± 0.01 | 54.52 ± 1.11 |

### 2.2 Experimental design

A pot experiment was performed in a greenhouse at the Graduate School of Hokkaido University. The sampled soil was air dried and sieved with 2 mm mesh and subsequently filled into Wagner pots (surface area = 1/5000 a). 1.8 kg of air dried soil was contained by each pot. The experimental design was completely randomized, included three fertilizer treatments × four mixed cropping treatments as well as was conducted in three replicates. The pots received one of the three types of fertilizer treatments, namely control (Ctr), chicken manure (50 g pot^− 1^ of carbonized chicken manure regarded as ‘CM’), or chemical fertilizer containing P and K (regarded as ‘CF’). The application rate for CF was 30 kg P ha^− 1^ and 50 kg K ha^− 1^. The soil as well as the CM chemical property were described in Table 1.

Afterwards, each pot received one of the four types of plant treatment: 1) single maize (SM), 2) the mixture of cowpea and maize (VM), 3) the mixture of velvet bean and maize (MM), or 4) the mixture of common bean and maize (PM). Three replications were included in each treatment. During the conduction of these treatments, maize (*Zea. Maize* L.) and three legume seeds, including cowpea (*Vigna unguiculate* (L.) Walp.), velvet bean (*Mucuna pruriens* (L.) DC.), and common bean (*Phaseolus vulgaris* L.) were sprouted for two weeks in small pots filled with vermiculite before transplantation to the Wagner pots. During the experiment, the temperature was maintained around 25°C to 30°C and the plants were grown for 50 days after the transplantation.

### 2.3 Chemical property analysis

The soils were sampled and measured for pH, extractable NH_4_^+^ and NO_3_^−^ concentrations 50 days after the transplantation. For soil pH, 6 g of soil was shaken for 30 min with 30 mL of Milli-Q water and pH was subsequently measured with a pH sensor (AS800, ASONE Co., Japan). For the extractable NH_4_^+^ and NO_3_^−^, the samples were extracted with KCl solution (2 mol L^− 1^) followed by colorimetric analysis using a flow injection analyzer system (ACLA-700, Aqualab Co., Ltd., Japan). Two-way analysis of variance (ANOVA) was afterwards performed to investigate the interaction between the environmental factors and the experimental treatments (Table 2).

**Table 2.**
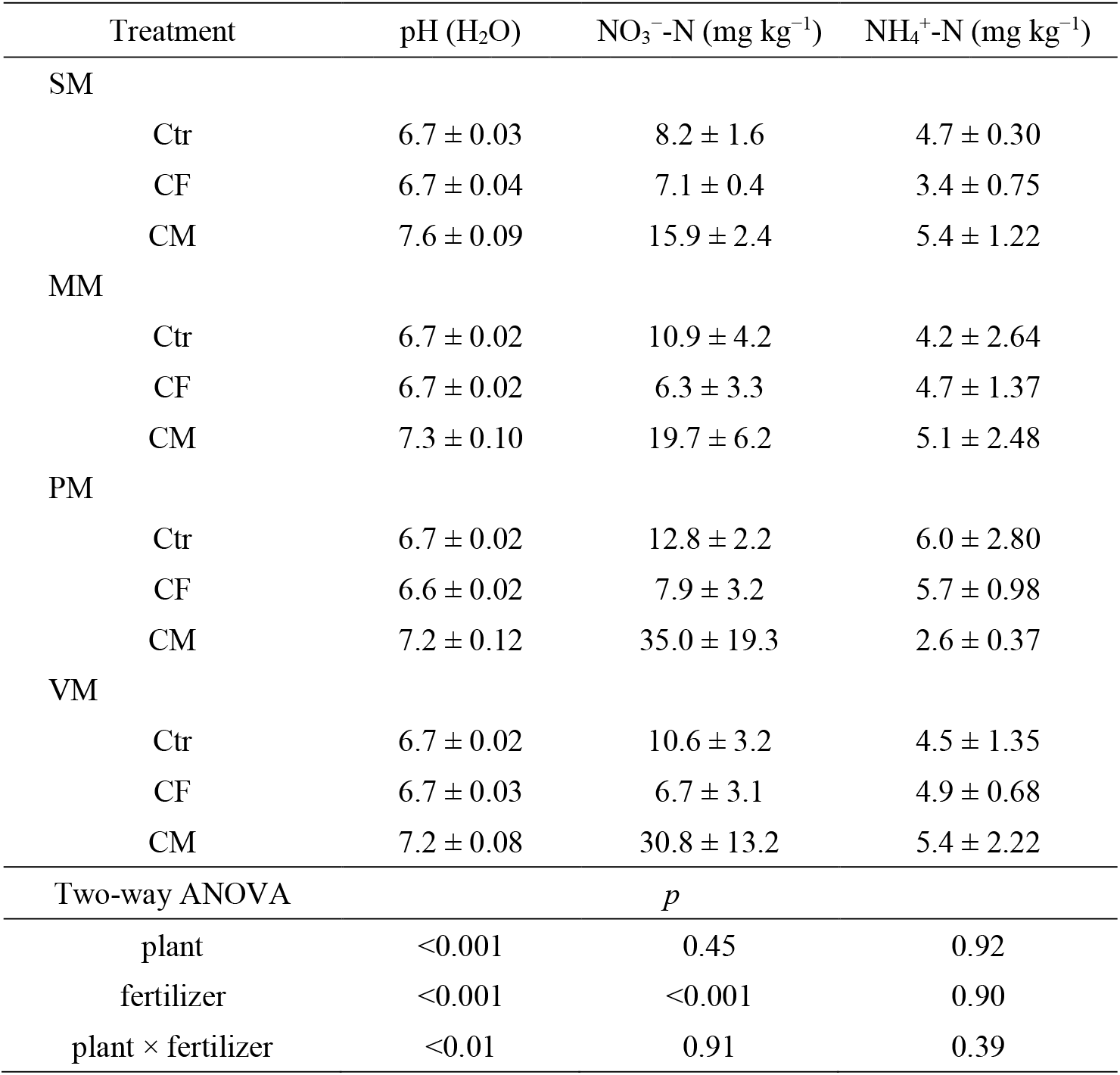
Soil pH, nitrate and ammonium content after plant cultivation. The expressed plant species included single maize, SM; maize cropped with velvet bean (*Mucuna pruriens* (L.) DC.), MM; maize cropped with common bean (*Phaseolus vulgaris* L.), PM; maize cropped with cowpea (*Vigna unguiculate* (L.) Walp.), VM. Fertilization was undertaken for the control, Ctr; chemical fertilizer, CF; and carbonized chicken manure, CM. Two-way ANOVA analysis was performed to examine the effects of interaction between the plant species and the fertilizer, and the p-values were shown at the bottom of the table.

| Treatment | pH (H<sub>2</sub>O) | NO<sub>3</sub><sup>-</sup>-N (mg kg<sup>-1</sup>) | NH<sub>4</sub><sup>+</sup>-N (mg kg<sup>-1</sup>) |
| --- | --- | --- | --- |
| SM |  |  |  |
| Ctr | 6.7 ± 0.03 | 8.2 ± 1.6 | 4.7 ± 0.30 |
| CF | 6.7 ± 0.04 | 7.1 ± 0.4 | 3.4 ± 0.75 |
| CM | 7.6 ± 0.09 | 15.9 ± 2.4 | 5.4 ± 1.22 |
| MM |  |  |  |
| Ctr | 6.7 ± 0.02 | 10.9 ± 4.2 | 4.2 ± 2.64 |
| CF | 6.7 ± 0.02 | 6.3 ± 3.3 | 4.7 ± 1.37 |
| CM | 7.3 ± 0.10 | 19.7 ± 6.2 | 5.1 ± 2.48 |
| PM |  |  |  |
| Ctr | 6.7 ± 0.02 | 12.8 ± 2.2 | 6.0 ± 2.80 |
| CF | 6.6 ± 0.02 | 7.9 ± 3.2 | 5.7 ± 0.98 |
| CM | 7.2 ± 0.12 | 35.0 ± 19.3 | 2.6 ± 0.37 |
| VM |  |  |  |
| Ctr | 6.7 ± 0.02 | 10.6 ± 3.2 | 4.5 ± 1.35 |
| CF | 6.7 ± 0.03 | 6.7 ± 3.1 | 4.9 ± 0.68 |
| CM | 7.2 ± 0.08 | 30.8 ± 13.2 | 5.4 ± 2.22 |
| Two-way ANOVA |  | *p* |  |
| plant | <0.001 | 0.45 | 0.92 |
| fertilizer | <0.001 | <0.001 | 0.90 |
| plant × fertilizer | <0.01 | 0.91 | 0.39 |

### 2.4 DNA extraction and 16S rRNA sequencing

Using the same sampled soils, DNA was extracted with NucleoSpin® Soil (Takara Bio Inc., Japan) according to the instructions of the manufacturer. The extracted DNA was subsequently amplified by polymerase chain reaction (PCR) targeting the V4 region of 16S rRNA (amplicon size ∼ 250 bp, forward primer = 515F: 5′-GTGCCAGCMGCCGCGGTAA-3′, reverse primer = 806R: 5′-GGACTACHVGGGTWTCTAAT-3′). To perform PCR, 10 µL AmpliTaq Gold® 360 Master Mix (Applied Biosystems, Foster City, CA, USA), 0.4 μL of the forward primer, 0.4 μL of the reverse primer, 7.2 μL nuclease-free water, and 1 μL of DNA extract were mixed. Then, the first PCR cycle was set to 95°C for 10 min, then 20 cycles at 95°C for 30 s, 57°C for 30 s, and 72°C for 1 min, followed by 72°C for 7 min. The PCR products were subsequently purified with Agencourt AMPure XP (Beckman Coulter) according to the given protocol. Then, the purified products were quantified with the QuantiFluor® ONE dsDNA System by a QuantusTM Fluorometer E6150 (Promega, Madison, USA).

Another PCR was performed with the utilisation of the amplicon-obtained products to make them Ion Torrent sequence sample-specific. To achieve this, the 515F forward primer with the Ion Xpress Barcode Adapters Kit sequence as well as the 806R reverse primer attached to the Ion Xpress sequence of the Ion P1 adaptor were used (Thermo Fisher Scientific K.K.). The first PCR products were diluted to 2000 ng mL^− 1^ and 1 μL of each product were subsequently mixed with 10 μL of AmpliTaq Gold® 360 Master Mix, 0.4 μL of the forward primer, 0.4 μL of the reverse primer, and 7.2 μL of nuclease-free water. The second PCR cycle was set to 95°C for 10 min, then 5 cycles at 95°C for 30 s, 57°C for 30 s, and 72°C for 1 min, followed by 72°C for 7 min. The second PCR products were purified in accordance with the same method outlined above. The final length and concentration of the amplicons were confirmed using a Bioanalyzer DNA 1000 Kit (Agilent Technologies, USA). The library was subsequently diluted to 50 pM and loaded into the Ion 318 chip using an Ion Chef Instruments with an Ion PGM Hi-Q Chef Solutions. Afterwards, samples were sequenced on an Ion PGM Sequencer with Ion PGM Hi-Q View Sequence Solutions (Ion Torrent Life Technologies, USA).

### 2.5 Sequence processing

The barcoded 16S rRNA gene sequences were denoised, quality-filtered, and assessed using the DADA2 algorithm implemented in Quantitative Insights Into Microbial Ecology (QIIME2) and with its workflow (Bolyen et al., 2019). Rarefaction was performed with minimal reads among all samples and sequence data were subsampled to 41095 sequences per sample. The R package Vegan (version 2.5.6) was used to access sample depth and generate the alpha rarefaction curve plot (Figure S1). The rarefaction curve was afterwards evaluated using the interval of step sample size, 1000.

### 2.6 Measurement of bacterial abundance

For the measurement of the bacterial abundance, quantitative PCR (qPCR) was performed using the extracted DNA, diluted 50 times with nuclease-free water. The 515F/806R primer pairs described above were used to amplify the V4 region of the 16S rRNA. For the standard curve, the PCR products from the DNA extracted from the control pots were used, which were purified with AMPure XP (Beckman Coulter) and further diluted to 5 stages of different concentrations. Samples were prepared with 10.4 μL of KAPA SYBR Fast qPCR kit (Kapa Biosystem, USA), 0.08 μL of the forward primer, 0.08 μL of the reverse primer, and 2 μL of diluted DNA extract. Moreover, nuclease-free water was added to achieve the final volume of 20 μL. CFX96 Touch™ Real (Bio-Rad Laboratories, Inc., USA) was used and the cycling condition was set to 95°C for 30 s, 35 cycles at 95°C for 30 s, 58°C for 30 s, and 72°C for 1 min, followed by 95°C for 1 min and subsequently 55°C to 95°C by 1°C increment for 10 s. The Ct values (Threshed cycle) were calculated after the quantification of the amplification results using version 1.4.1 of the qR R package.

### 2.7 Statistical analysis

To quantify the diversity of soil microbial communities, the Shannon index (Shannon, 1948) and Simpson index (Simpson, 1949), an estimation of community alpha diversity, were calculated. For each diversity index, two-way analysis of variance (ANOVA) was performed using fertilizer treatments and plant species treatments as factors with the emmeans R package, version 1.4.7 (Lenth et al., 2020). Multiple comparisons were subsequently performed using the Turkey–Kramer method.

Regarding the phylum level of beta diversity (the difference between samples), non-metric multidimensional scaling based on the Bray–Curtis distance metrics was used to visualize sample dissimilarities. Permutational analysis of variance (PERMANOVA) (permutation = 9999) was conducted and significant phyla (p < 0.001) were subsequently identified for the interactions between the fertilizer and the plant type treatments, using two-way ANOVA.

Unique operational taxonomy units (OTUs) for each plant treatment within the same fertilizer treatment were extracted using Venn diagrams. Version 1.6.20 of the VennDiagram R package was applied using the co-occurred OTUs among the three replications for each treatment. The relative abundance of unique OTUs within the total abundance was plotted for the treatments with significant differences in diversity indices. Similarly, the top 20 most abundant OTUs on the average for all samples were extracted to estimate the effects of each treatment on the relative abundance of the dominant OTUs. The relative abundance of extracted OTUs was illustrated with the class level of classification.

## 3 Results

### 3.1 Diversity indices and abundance of bacterial communities

The interaction effect between plant species and fertilization was shown by soil microbial diversity indices. Within the PM treatment, a significantly lower Simpson’s diversity indicator was observed with no fertilization (Ctr) when compared to the treatment with CM (Figure 1). In contrast, relatively lower diversity was shown by the MM treatment with the CM application compared to the Ctr treatment.

**Figure 1.**
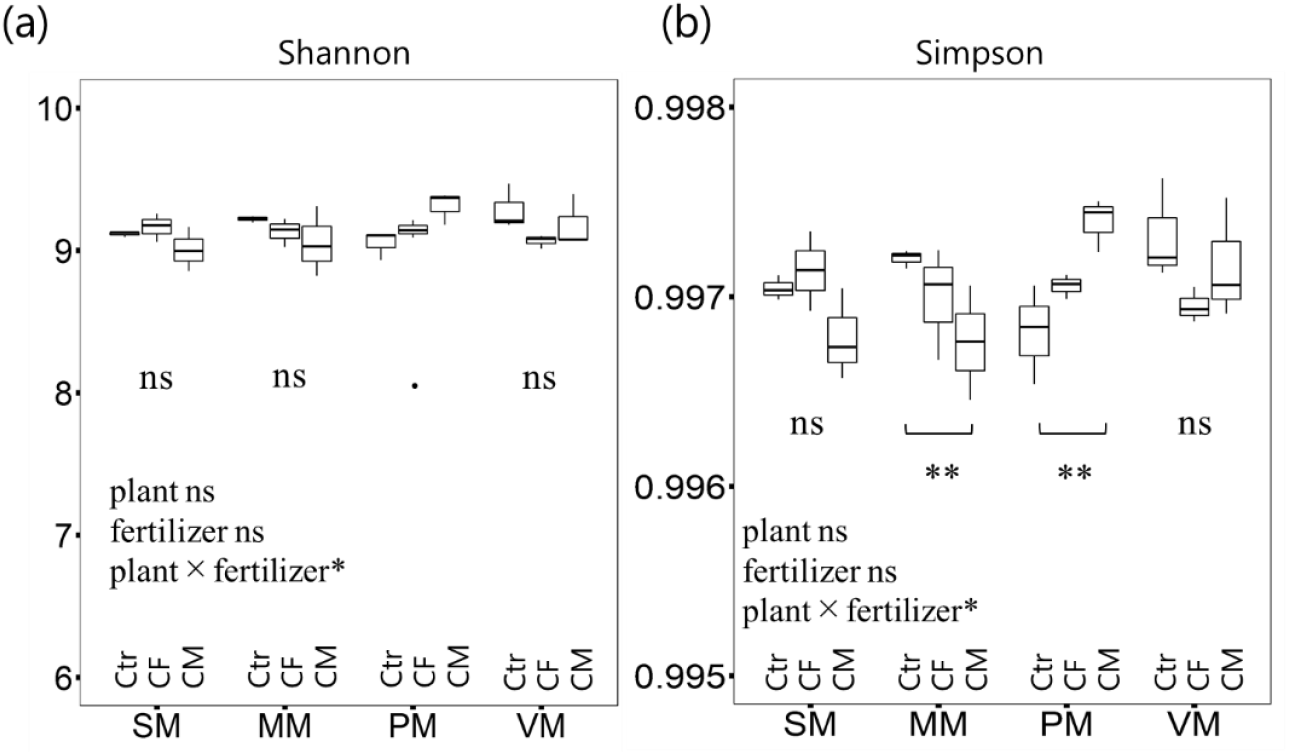
The box plot of bacterial OTUs alpha diversity indicating (a) the Shannon’s Index and (b) the Simpson’s index. Two-way ANOVA and Tukey–Kramer’s pairwise comparison were performed on the calculated alpha-diversity indices: **·** p < 0.25, * p < 0.05, ** p < 0.01. The abbreviations of plant species were as follows: single maize, SM; maize cropped with velvet bean, MM; maize cropped with common bean, MP; maize cropped with cowpea, VM. Fertilizer treatments were abbreviated as no fertilizer, Ctr; chemical fertilizer, CF; and carbonized chicken manure, CM.

The bacterial absolute abundance, based on the qPCR analyses, showed a significant impact of plant treatments when averaged across fertilizer types (Table S1). The bacterial abundance of the SM and VM treatments was significantly different. However, there was no correlation between bacterial absolute abundances and diversity indicators.

The analysis of the soil 16S rRNA gene sequence provided 738 OTUs to 1196 OTUs per sample. The Venn diagrams showed that soils under the Ctr, CF, and CM treatment had 297, 293, and 283 core OTUs (the overlapped area across plant treatments), respectively (Figure 2) as well as increasing numbers of unique OTUs with increasing diversity indices, for example, the unique OTU number increased from 28 to 84 for PM in case of the comparison of Ctr and CM, whereas it decreased from 52 to 35 in MM. The relative abundance of these unique bacteria within the whole bacterial abundance increased from 1.9% (Ctr) to 5.2% (CM) in the PM but decreased from 4.3% (Ctr) to 1.9% (CM) in the MM treatment (Figure 3). The abundance of Acidobacteria and Verrucomicrobia was suggested by the community structures of these unique bacterial taxa to dominate within the unique microbial communities and was correlated to the increase of unique OTU numbers during the comparison of different treatments. In contrast, the abundance of Gemmatimonadates, Chloroflexi and Planctomycetes was not correlated to the increase in the unique OTU numbers, although they were considered to be dominant phylum within the unique OTUs.

**Figure 2.**
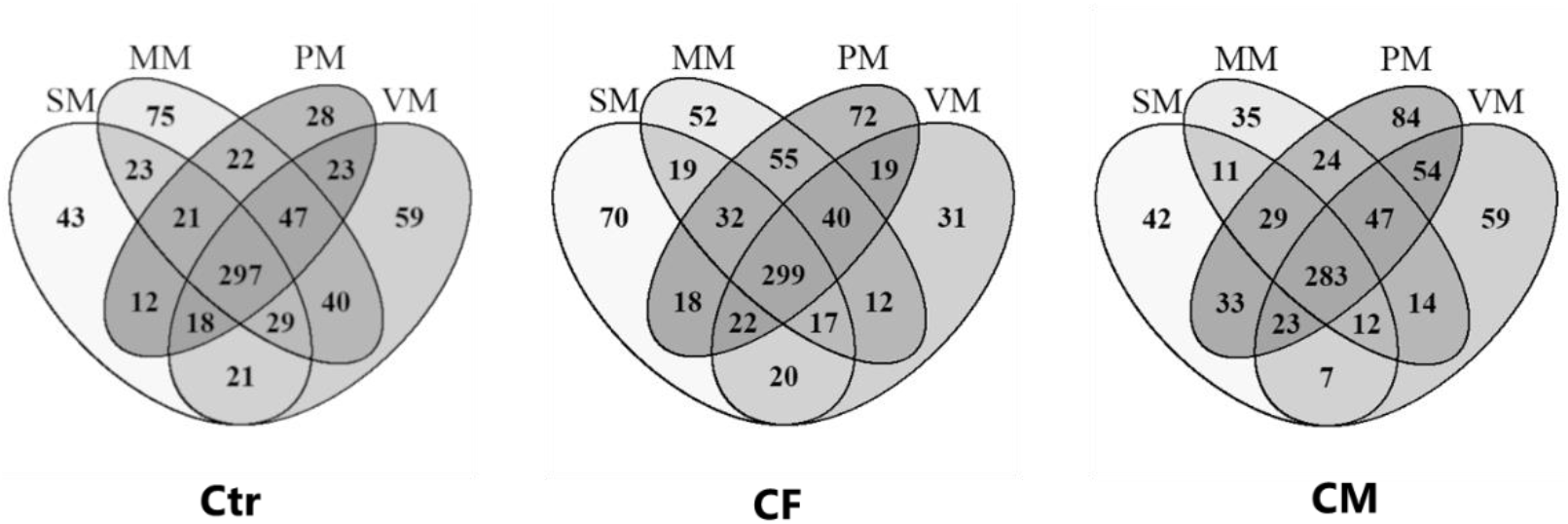
The distribution of OTUs among different mixed cropping systems within the fertilized treatment. The number showed the shared and unique OTUs in each treatment. Overlapping circles showed the shared OTUs among the treatments. Each circle included OTUs co-occurred among three replications.

**Figure 3.**
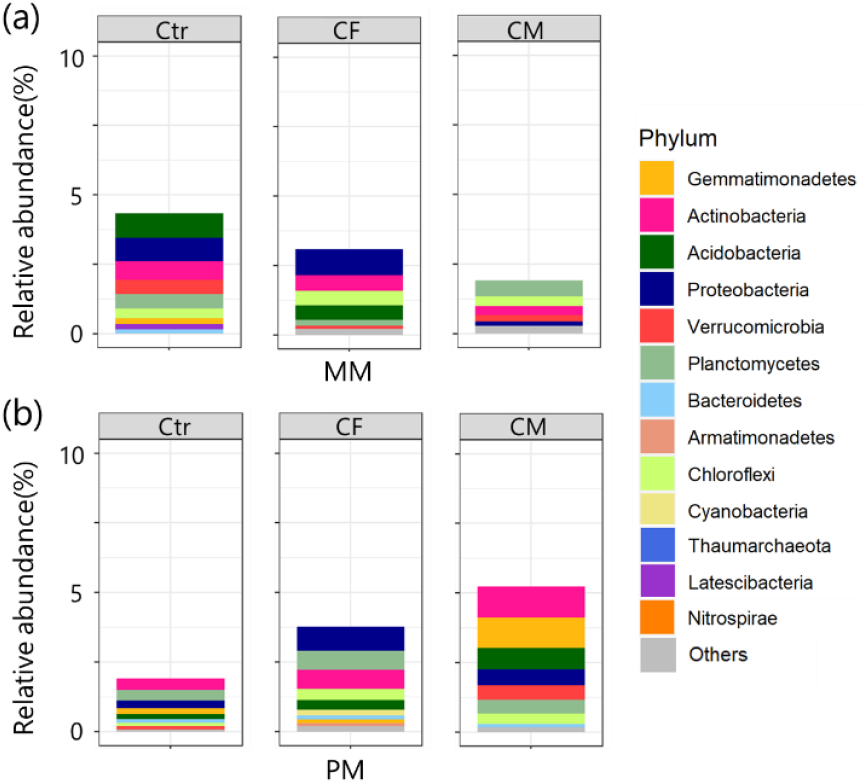
The relative abundance of unique OTU phyla found in MM (a) and PM (b) treatments. The bar plots indicated the total relative abundance dimension with averaged value across the replications. Only phyla with a relative abundance > 0.1% in averaged value were represented.

### 3.2 Soil chemical properties

Positive impacts were shown by the soil pH of the CM application for all the plant treatments (Table 2). There was no interaction effect observed on the concentration of soil NO_3_^−^-N and NH_4_^+^-N, but the effect of fertilizer was perceived on the NO_3_^−^-N concentration (p < 0.001). With the CM application, the NO_3_^−^-N concentration was significantly increased (p < 0.05).

### 3.3 Top 20 dominant OTUs analysis

Within the top 20 dominant OTUs, the CM application increased the relative abundance of the Soil Crenarchaeotic Group (SCG) belonging to Thaumarchaeota as well as decreased the relative abundance of Alphaproteobacteria for the treatments excluding the PM (Figure 4). In the PM treatment, the abundance of SCG decreased when CM was applied in comparison with the Ctr. Furthermore, when the cumulative abundance of the top 20 dominant OTUs was compared between the Ctr and the CM, a decrease was observed from 15.8% (Ctr) to 14.1% (CM) in the PM treatment but an increase from 14.9% (Ctr) to 16.3% (CM) was identified in case of the MM treatment.

**Figure 4.**
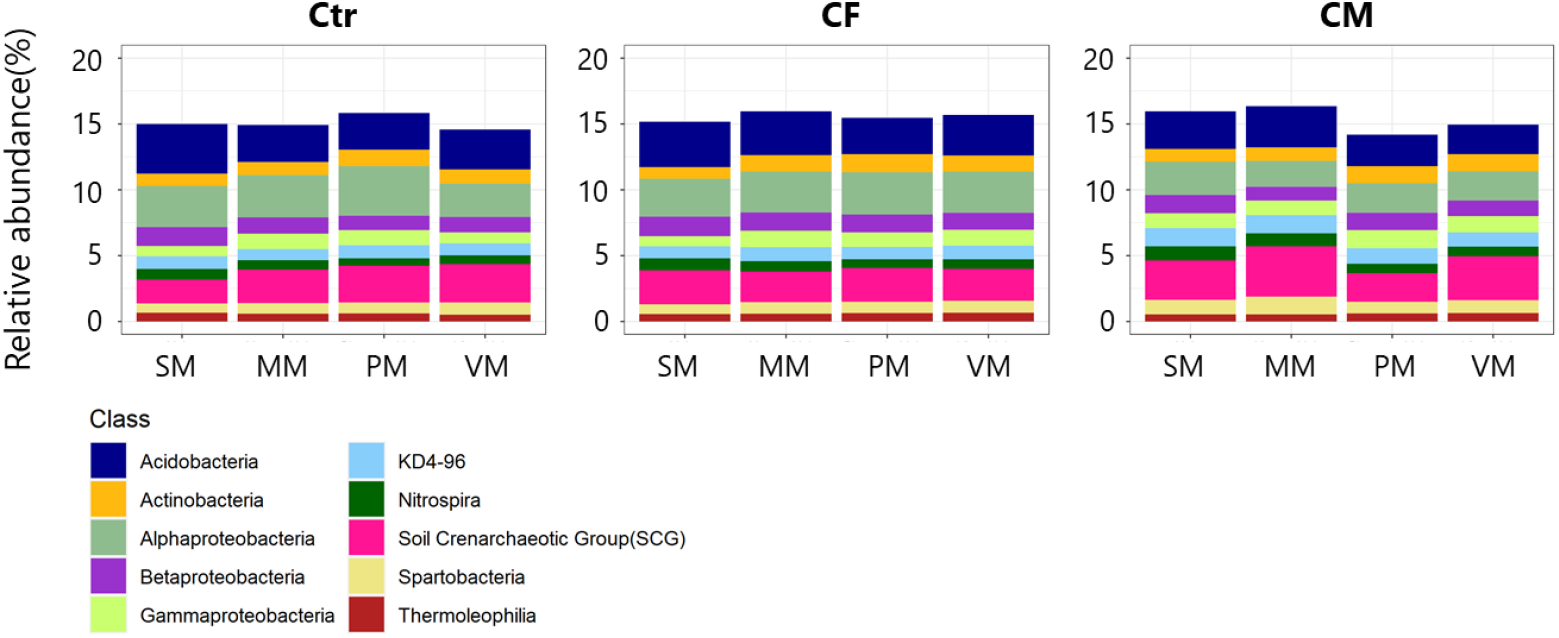
Relative abundance of the top 20 most abundant OTUs with each bacterial class. The top 20 OTUs were extracted based on the average of all samples and their relative abundance was visualized with class level classification.

### 3.4 Environmental factors and dominant OTUs composition

The phylum level of the bacterial communities was significantly influenced by fertilizer treatments (p < 0.05). The impacts of environmental factors on the community structures were p = 0.593, 0.118 and < 0.001 for NH_4_^+^-N, NO_3_^−^-N and pH, respectively. The relative abundance of Thaumarchaeota, Armatimonadetes, Chloroflexi, Planctomycetes Verrucomicrobia, and Proteobacteria contributed to the changes in the community structures (Figure 5) as well as their abundance was influenced by the interaction between the plant and fertilizer treatments (Table 3). Within the CM treatment, Thaumarchaeota, Chloroflexi, Planctomycetes, and Verrucomicrobia were significantly higher in the MM treatment than the PM treatment (Table 4).

**Table 3.** Bacteria phylum with permutational p-value < 0.001 (permutation = 9999). Two-way ANOVA with the plant species and the fertilizer treatment as factors was performed and the p-values were shown in the table.

| Phylum | Two-way ANOVA |  |  |
| --- | --- | --- | --- |
|  | fertilizer | plant | plant ×<br>fertilizer |
| *Thaumarchaeota* | 0.018 | 0.19 | <0.01 |
| *Acidobacteria* | <0.001 | <0.01 | 0.063 |
| *Actinobacteria* | 0.35 | <0.01 | 0.074 |
| *Armatimonadetes* | <0.001 | 0.0021 | <0.01 |
| *Bacteroidetes* | 0.097 | 0.5 | 0.12 |
| *Chloroflexi* | <0.001 | <0.001 | <0.001 |
| *Gemmatimonadetes* | 0.48 | 0.51 | 0.048 |
| *Latescibacteria* | 0.34 | 0.18 | 0.48 |
| *Nitrospirae* | <0.001 | <0.001 | 0.47 |
| *Planctomycetes* | <0.001 | 0.096 | <0.001 |
| *Proteobacteria* | <0.001 | 0.16 | <0.01 |
| *Verrucomicrobia* | <0.001 | <0.001 | <0.001 |

**Table 4.** The relative abundance of the phylum showed a significant interaction between the plant species and the fertilizer treatment. The results from multiple pairwise comparisons shown as different alphabets and different letters indicated significant differences between the treatments (p < 0.05).

| Treatment | Relative abundance (%) |  |  |  |  |  |
| --- | --- | --- | --- | --- | --- | --- |
|  | *Thaumarchaeota* | *Armatimonadetes* | *Chloroflexi* | *Planctomycetes* | *Verrucomicrobia* | *Proteobacteria* |
| Ctr |  |  |  |  |  |  |
| SM | 3.23 a | 0.58 a | 7.84 ab | 6.76 a | 2.93 a | 29.93 b |
| MM | 4.70 ab | 0.46 a | 7.58 a | 8.53 b | 4.37 b | 28.19 ab |
| PM | 5.22 ab | 0.42 a | 7.84 ab | 8.21 ab | 4.20 b | 27.92 ab |
| VM | 5.66 b | 0.53 a | 8.74 b | 10.74 c | 4.57 b | 25.13 a |
| CF |  |  |  |  |  |  |
| SM | 4.94 a | 0.62 a | 8.76 b | 9.75 b | 4.08 a | 26.95 a |
| MM | 4.21 a | 0.60 a | 8.03 ab | 8.48 ab | 4.43 a | 26.49 a |
| PM | 4.72 a | 0.57 a | 7.95 ab | 8.70 ab | 3.93 a | 27.05 a |
| VM | 4.23 a | 0.53 a | 7.71 a | 7.63 a | 3.78 a | 28.31 a |
| CM |  |  |  |  |  |  |
| SM | 5.50 ab | 0.83 b | 12.10 c | 10.94 b | 5.89 b | 22.79 ab |
| MM | 6.67 b | 0.72 b | 12.27 c | 11.19 b | 6.19 b | 20.04 a |
| PM | 4.01 a | 0.76 b | 10.65 b | 9.09 a | 4.56 a | 24.21 b |
| VM | 6.19 b | 0.48 a | 8.70 a | 7.94 a | 4.50 a | 25.14 b |

**Figure 5.**
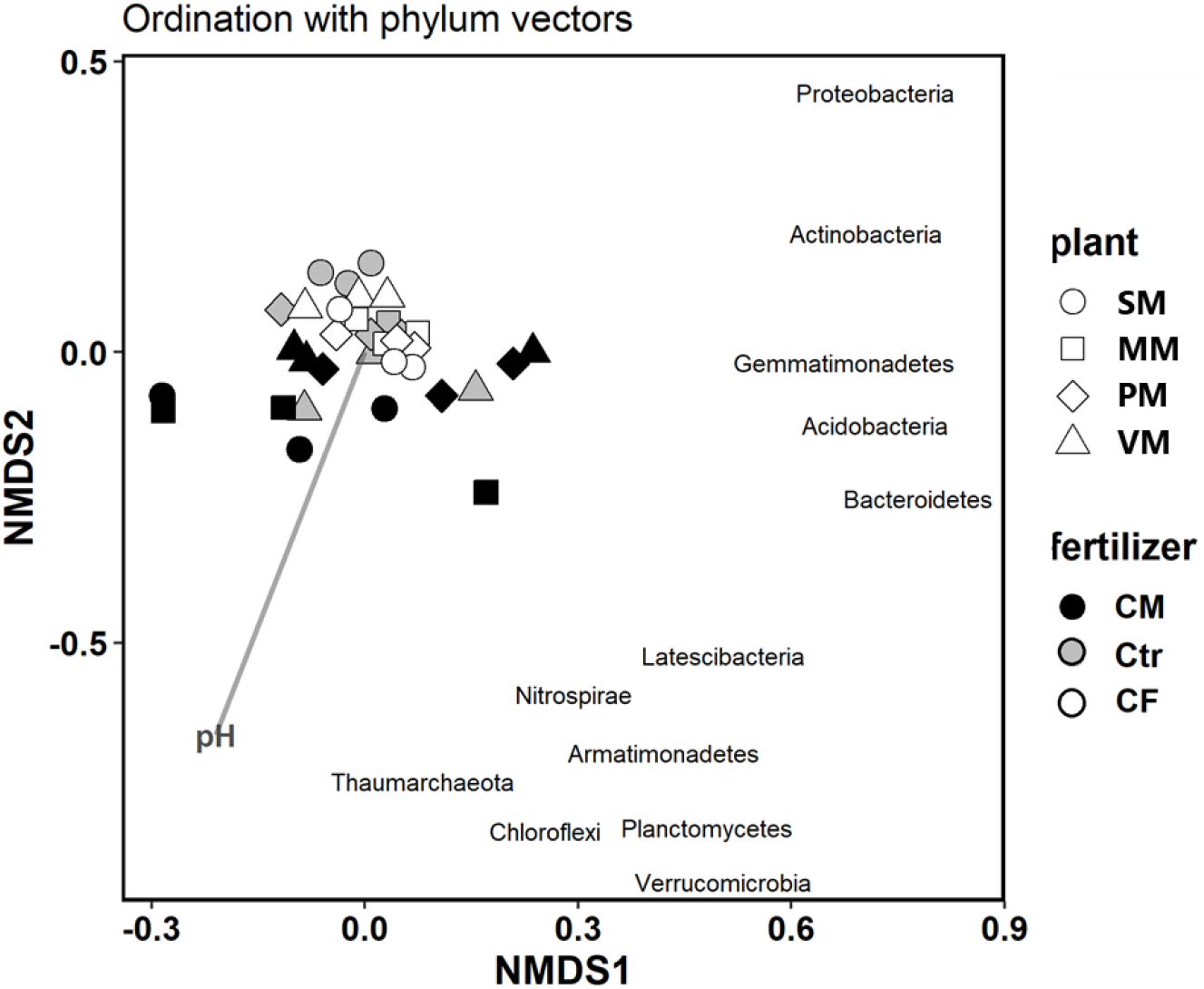
Non-metric multidimension scaling plot based on Bray–Curtis dissimilarity with significant bacterial phylum (p < 0.001) and environmental factors (stress 0.131). The bacterial community was significantly influenced by fertilizer treatment (p < 0.05) and pH (p < 0.001).

## 4 Discussion

### 4.1 Characteristics of unique bacteria in each treatment

Our experimental results demonstrated that mixed cropping systems diversified soil microbial communities only in case of the use of specific fertilizer and legume species (Figure 1) as well as there was no universal impact of the mixed cropping on the soil bacterial diversities identified. No consistent statement about microbial diversity in multiple cropping systems using cereal and legume was confirmed by previous studies. For example, Fu et al. (2019) and Kihara et al. (2012) found that an intercropping system with maize and soybean increased microbial diversity, whereas Yu et al. (2019) showed that there was no significant impact of the intercropping system on microbial diversity. Moreover, a complex interaction with N fertilization regarding the impact of intercropping on microbial diversification was shown by Zhang et al. (2020). Therefore, as demonstrated in this study, both plant species and fertilizer types were found to be related to the mechanisms behind the diversification of soil microbes and our study highlighted the importance of the use of CM, when compared to chemical fertilizer, in association with the diversification of soil microbes.

In this experiment, the number and abundance of unique OTUs were found to be in control of the magnitudes of the microbial diversities, particularly when CM treatments were compared with the Ctr (Figure 1, Figure 2). To support this result, the unique OTU numbers were also shown by previous studies to correspond to soil microbial diversity indices (Deng et al., 2020; He et al., 2019). In regard of unique OTUs accounting for a small proportion of the total community, however, Chen et al. (2020) and Jousset et al. (2017) claimed that these small and rare microbial taxa (less than 5%) should be recognized as the vulnerable components in association with the diversity of functionality in agricultural ecosystems. Therefore, it is valuable to focus on communities with unique OTUs to elucidate the microbial species driving the community diversity and the functional variabilities.

Interestingly, the unique bacterial abundance and the diversity in the PM were increased by the addition of CM, whereas the CM decreased the abundance and diversity in MM. Thus, the plant species were identified to be an important factor controlling the presence of the unique bacteria when CM was applied as different plants have different extent of ability to recruit specific bacterial communities (Zhang et al., 2019). It is possible that the amount and quality of root exudates were differently impacted when the PM and MM were compared as soil microbes are known to show a preference in association with the exudate (substrates) supplied from the roots (Duchene et al., 2017; Zhalnina et al., 2018). Our study indicated that both the legume species and the fertilizer types should be considered for soil microbial diversity conservation in agricultural ecosystems. Further studies are needed to investigate the chemical composition of the root exudates and their impact on the unique bacterial community as well as the interaction with the use of organic fertilizers.

In regard of the unique OTUs, the abundance of Acidobacteria and Verrucomicrobia was positively correlated with the % of unique OTUs and the diversity indices (Figure 3). One of the reasons for this positive correlation can be that it is due to Acidobacteria being one of the dominant bacteria in the rhizosphere (Barns et al., 2007). In rhizosphere soils, these bacteria often dominate the present bacterial communities due to their relatively higher metabolic potential (Lee et al., 2008). Moreover, whole-genome sequence analyses of some Acidobacteria strains demonstrated that they are related to carbohydrate metabolism and utilise root exudates (de Chaves et al., 2019; Kielak et al., 2016; Nielsen et al., 2014). In addition, these bacteria have functional genes in association with N cycles, including NO_3_^−^-N and NO_2_^−^-N reduction (Ward et al., 2009). Furthermore, Verrucomicrobia were also reported to hold diverse communities in the rhizosphere (Bünger et al., 2020) and their role in polysaccharide degradation processes and methane metabolization were identified (Dunfield et al., 2007; He et al., 2017; Pold et al., 2018). Additionally, Verrucomicrobia were observed to be involved in N cycling processes, such as N fixation and ammonia oxidation (Cabello-Yeves et al., 2017; Mohammadi et al., 2017; Nixon et al., 2019). Thus, our results suggest that root-associated bacteria uniquely associated with the use of specific fertilizer and legume species control C and N cycles in soils to some extent. However, limited research was performed to elucidate the functional role of unique OTUs in the agricultural ecosystem; therefore this remains an aim for future investigations.

### 4.2 Characteristics of the top twenty most abundant OTUs

Regarding the top twenty most abundant OTUs in the PM treatment, the abundance of SCG that belongs to the Thaumarchaeota and Alphaproteobacteria conspicuously decreased when the CM treatment was compared to the Ctr treatment (Figure 4). Generally, the community structures of bacteria were found to fluctuate due to changes in soil pH, salt-based ions, including potassium, sodium, and calcium ions (Wang et al., 2020). The CM treatment altered these soil chemical characteristics in the current experiment. However, it is difficult to clearly explain why the response of some bacterial OTUs to the CM application depended on the legume species, thus this remains a question to be answered in future studies.

It is important to highlight that the PM with CM application was characterized by the relatively lower abundance of the top 20 OTUs, but the relatively higher abundance of the rare OTUs, compared to other legume species with CM application, as discussed earlier. Therefore, the specific combinations of plant species and fertilizer (e.g. PM with CM) were found to potentially inhibit dominant bacterial growth while enhancing the number and the abundance of unique bacteria. The conflict between dominant and unique bacterial growth might be related to microbial diversity differences among the different legume-based mixed cropping systems.

### 4.3 Effects on other bacterial phyla

During the comparison of the community structures at the phylum level within the CM applied soils, the abundance of Thaumarchaeota, Chloroflexi, Planctomycetes and Verrucomicrobia was significantly lower in the PM in comparison with other legume species (Table 4). Thaumarchaeota is known as ammonia-oxidizing archaea ubiquitously present in a wide variety of ecosystems (Brochier-Armanet et al., 2012; Pester et al., 2011; Spang et al., 2010). This phylum was reported to prefer to live in low NO_3_^−^-N concentration (Oton et al., 2016). Thus, Thaumarchaeota abundance might be decreased only in PM with CM application due to the relatively higher NO_3_^−^-N concentrations in this treatment (Table 2). Similarly, the phylum of Chloroflexi is known to be sensitive to the amount of NO_3_^−^-N, reducing their abundance with increasing NO_3_^−^-N levels (Lou et al., 2019; Z. Zhang et al., 2020). Furthermore, Planctomycetes were reported to be negatively influenced by the increasing amount of NO_3_^−^-N (Blanchart et al., 2006). In addition, Verrucomicrobia are known to be abundant in grasslands and at subsurface soil horizons (Bergmann et al., 2011). Their abundance is strongly regulated by soil pH and the soil C/N ratio (Aguirre-von-Wobeser et al., 2018; Shen et al., 2017). However, limited functionality and information on habitat are known as Verrucomicrobia is difficult to be isolated and cultured.

It is important to note that CM treatment did not only increase soil NO_3_^−^-N levels, but the C availability in soils as well. Besides, previous studies often reported that the phylum mentioned in the previous paragraphs was influenced rather positively by the C availability. For example, the abundance of Thaumarchaeota, Chloroflexi and Planctomycetes was reported to have positive correlations with increasing soil organic C (Krzmarzick et al., 2012; Oton et al., 2016; Tao et al., 2017; Yan et al., 2019). Thus, further studies are needed to understand how microbial taxa respond in case of the presence of both positive and negative factors in various types of soil, including the increase in organic C and NO_3_^−^-N, as observed in our study, as the sensitivity of the microbial taxa to these factors might be important regarding the maintenance of soil microbial diversities.

## 5 Conclusion

It was demonstrated that mixed cropping systems can be useful to diversify soil bacterial communities only under a specific combination of legume species and fertilizer. The soil bacterial diversity was increased when CM was used for the mixed cropping of common bean and maize. In this treatment, the number of unique OTUs (treatment specific OTUs) was increased with increasing bacterial diversity indices. In addition, the unique OTUs also correlated with the abundance of root-associated bacteria, including Acidobacteria and Verrucomicrobia. In contrast, the relative abundance of the top 20 most abundant OTUs was inhibited in diversified bacterial communities. These phenomena (the increase of unique bacteria and the suppression of dominant OTUs) can be a reason for the perceived increases in diversity indices with the use of CM under the mixed cropping of common bean and maize. Further studies are needed to understand the functional changes in soils in association with the diversification of soil bacterial communities under the use of various legume species and fertilizers, particularly focusing on the unique bacterial communities.

## Supplementary

**Table S1.** The 16S rRNA gene abundance. The number of replications was 3 for each treatment. Two-way ANOVA was performed to examine the effect of interaction between the plant species and the fertilizer, and the results for the multiple pairwise comparisons were shown with different letters.

| Treatment | Log (copy number) g soil<sup>-1</sup> |  |
| --- | --- | --- |
| SM |  |  |
| Ctr | 12.2 ± 0.0 |  |
| CF | 12.2 ± 0.1 | b |
| CM | 12.2 ± 0.1 |  |
| MM |  |  |
| Ctr | 11.8 ± 0.0 |  |
| CF | 12.2 ± 0.1 | ab |
| CM | 11.9 ± 0.2 |  |
| PM |  |  |
| Ctr | 12.2 ± 0.2 |  |
| CF | 12.1 ± 0.1 | ab |
| CM | 12.0 ± 0.1 |  |
| VM |  |  |
| Ctr | 11.7 ± 0.1 |  |
| CF | 12.0 ± 0.2 | a |
| CM | 12.1 ± 0.1 |  |
| Two-way ANOVA | *p* |  |
| plant | <0.01 |  |
| fertilizer | 0.045 |  |
| plant × fertilizer | 0.069 |  |

**Figure S1.**
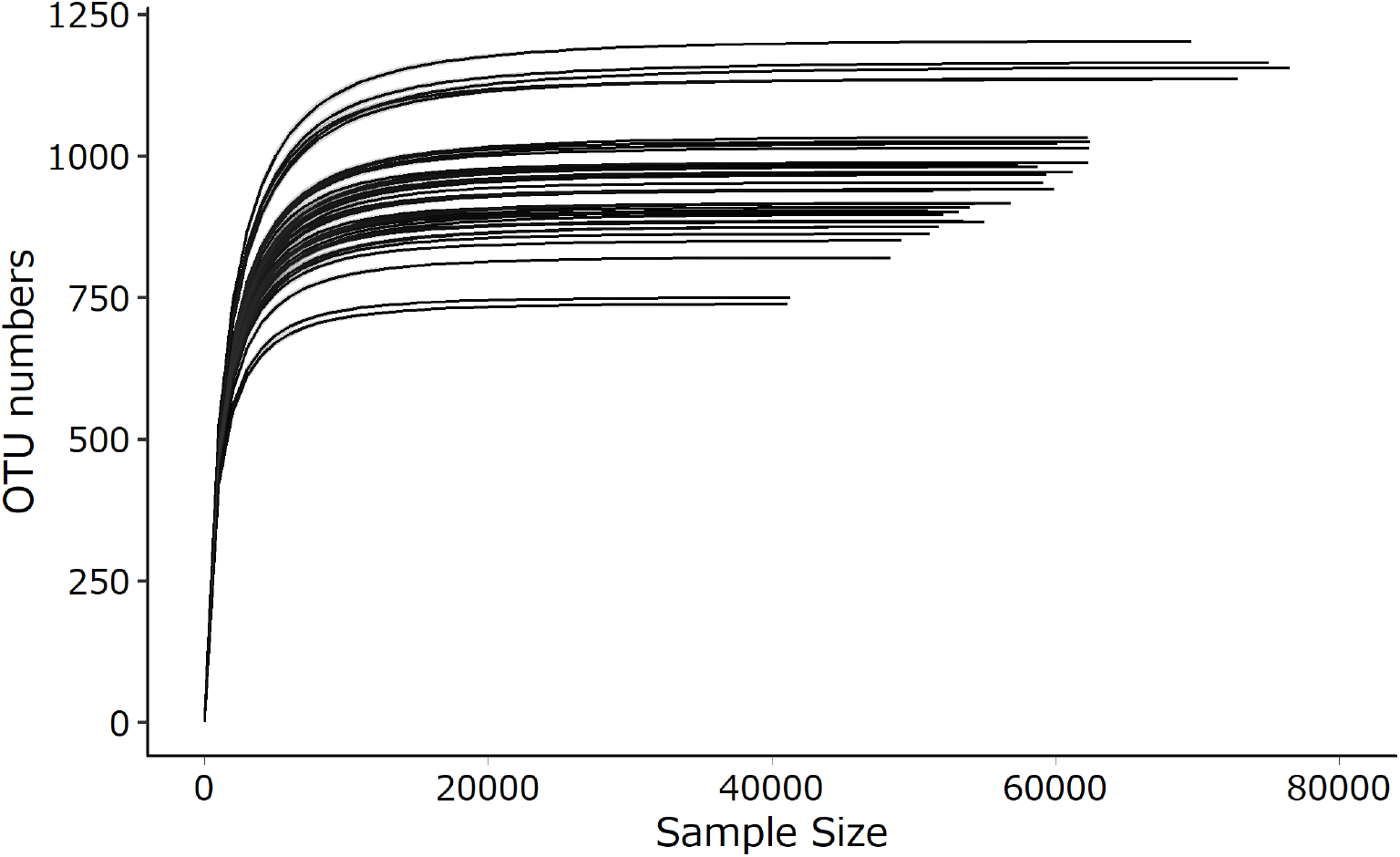
Alpha rarefaction curve of all samples. The curves were evaluated using the interval of step sample size, 1000.

**Figure S2.**
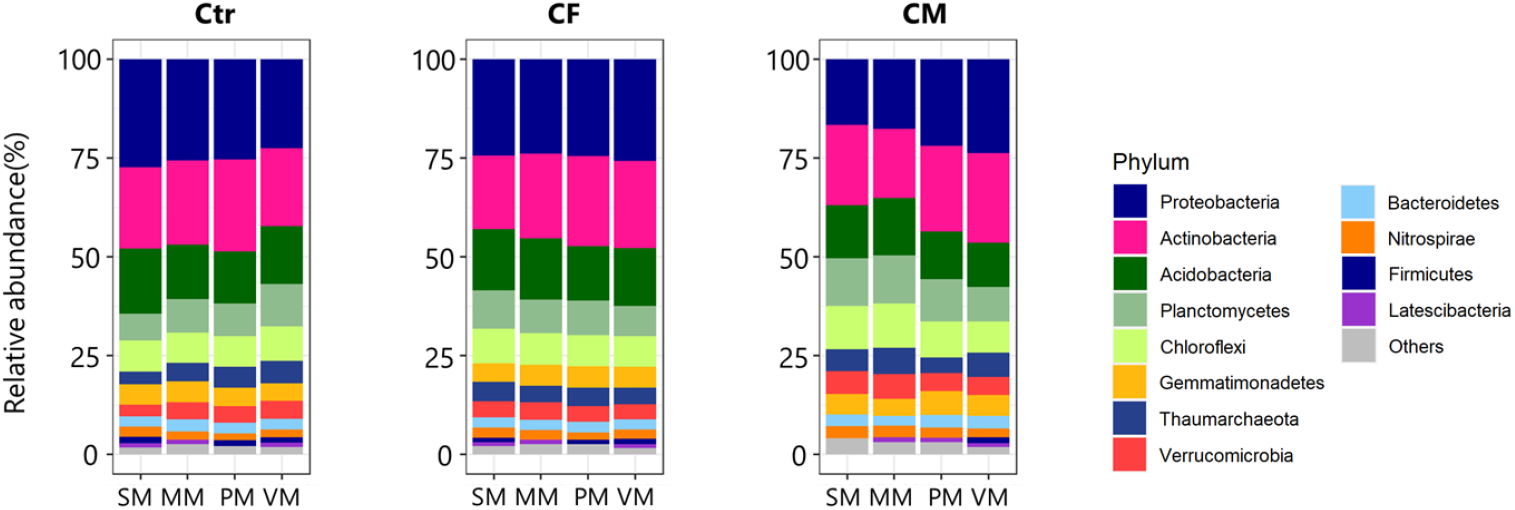
Relative abundance of the total community based on phylum-level classification. Only phyla with a relative abundance >0.1% in averaged value were represented. Plant treatments were shown as single maize, SM; velvet bean + maize, MM; common bean + maize, PM; and cowpea + maize, VM. Fertilizer treatments were shown as control, Ctr; chemical fertilizer, CF; and carbonized chicken manure, CM.

